## Supporting Appendix 1 for "A hydro-osmotic coarsening theory of biological cavity formation"

### 1 Derivation of the model for two lumens

#### 1.1 Lumen equations

Below we detail the conservation equations in the lumens and bridge following the the notations of Fig. 1D in the main text.

**Force balance** Force balance in a lumen $i$ is given by (i) the tension balance at the junction between the lumen and the bridge, which for an opening angle $\theta$ reads

$$\cos\theta = \frac{\gamma_c}{2\gamma}, \tag{1}$$

where $\gamma$ is the lumen tension, $\gamma_c$ is the contact (bridge) tension, and (ii) the Laplace's law

$$\delta P_i \equiv P_i - p_0 = \frac{\gamma}{R_i} = \frac{\gamma\sin\theta}{L_i}$$

with $R_i = \frac{L_i}{\sin\theta}$ the radius of curvature of the lumen.

**Mass balance** In 2D, the mass corresponds to the area of the lumen $A_i = \frac{L_i^2}{\mu}$, where $\mu = \frac{\sin^2\theta}{2\theta - \sin(2\theta)}$. Mass balance reads

$$\frac{dA_i}{dt} = 4\theta R_i \lambda_v \left[\delta\Pi_i - \delta P_i\right] - J_i^v, \tag{2}$$

where $\delta\Pi_i = \mathcal{R}T(C_i - c_0) \equiv \mathcal{R}T\delta C_i$, is the osmotic pressure jump (or concentration jump $C_i = \frac{N_i}{A_i}$) across the membrane and $J_i^v$ the longitudinal solvent flux toward the bridge. Using the geometrical relationship $L_i = \sqrt{\mu A_i} = R_i \sin\theta$, we can rewrite the equation above as a function of $L_i$ :

$$\frac{dL_i}{dt} = 2\mu\nu\lambda_v \left[\mathcal{R}T\delta C_i - \frac{\gamma\sin\theta}{L_i}\right] - \frac{\mu}{2L_i}J_i^v, \tag{3}$$

where we introduced $\nu = \frac{\theta}{\sin\theta}$. Non-dimensionalization of this equation leads to Eq.[6] in the main text.

Assuming no flux $J_i^v$, this mass balance equation tells us that at steady-state, the stationary lumen volume is given by $\delta\Pi_i = \delta P_i$, leading to $\frac{\delta C_i}{c_0} = \frac{\gamma}{\Pi_0 R_i} \preceq 10^{-2} \ll 1$, where we took $R_i \succeq 100\text{nm}$, $\gamma \sim 100\text{pN}/\mu\text{m}$ and $\Pi_0 \sim 10^5\text{Pa}$ (see Supporting Table 1). Therefore in the following, we will assume that the concentration jump is small compared to the cellular concentration $\delta C_i \ll c_0$.

**Solute balance** Similarly, the change in solute number $N_i$ is due to passive exchanges: across the membrane, driven by chemical potential gradients $\mu = \mathcal{R}T\log(c)$, and toward the bridge via a longitudinal flux $J_i^s$. An active pumping rate $j^a$ is added at membranes, yielding

$$\frac{dN_i}{dt} = 4\theta R_i \left[\lambda_s \mathcal{R}T \log\frac{c_0}{C_i} + j^a\right] - J_i^s, \tag{4}$$

Using $\delta C_i \ll c_0$, we can expand the log as $\log\frac{1}{1+\delta C_i/c_0} \underset{0}{\sim} -\frac{\delta C_i}{c_0}$ yielding

$$\frac{dN_i}{dt} = 2\nu L_i \left[-\lambda_s \mathcal{R}T\frac{\delta C_i}{c_0} + j^a\right] - J_i^s, \tag{5}$$

Non-dimensionalization of this equation will lead to Eq [7] in the main text.

#### 1.2 Bridge equations

The bridge is modeled as a 2-dimensional tube of diameter $e(x,t)$, of length $\ell(t)$. In 3D-geometry, the bridge is not a tube, rather a 2D sheet. We choose to parametrize the x-axis such that $x \in [-\frac{\ell}{2}, \frac{\ell}{2}]$ for symmetry reasons. Hydrostatic pressure jump between the bridge and the exterior is denoted $\delta p(x,t) = p(x,t) - p_0$, concentration jump $\delta c(x,t) = c(x,t) - c_0$.

**Force balance** The membranes have a tension $\gamma_c$, and adhesive molecule (cadherins) are distributed along the bridge with line density $\rho(x,t)$, attached to the two membranes, creating an elastic restoring force if the membranes are moved apart. Adding the contribution of the membrane tension, force balance reads :

$$k\rho(x,t)(e(x,t) - e_0) - \gamma_c \nabla^2 e(x,t) = \delta p(x,t), \tag{6}$$

where $k$ is the stiffness of individual cadherins molecules, $e_0$ is the reference distance between the two membranes, i.e. the bridge diameter.

In the following, we assume that the density and stiffness of cadherin molecules is large enough to ensure a very small deformation of the bridge $e(x,t) \sim e_0$, while keeping the pressure in the bridge $\delta p(x,t)$ of order unity. Therefore, we will neglect spatio-temporal variations in bridge thickness in the next.

**Mass balance** The local mass balance in the bridge is given by

$$(e(x+dx,t)dx - e(x,t))dx = [q(x,t) - q(x+dx,t)]\, dt + 2j_v.dx.dt \left[1 + \frac{1}{2}\left(\frac{\partial e(x,t)}{\partial x}\right)^2\right],$$

where $j_v$ is a lateral incoming flux, $q(x,t)$ is the longitudinal hydrodynamic flow along the x-axis, given by 2D Poiseuille flow:

$$q(x,t) = -\kappa_v \frac{\partial \delta p}{\partial x},$$

with $\kappa_v$ the effective friction, such that $\kappa_v = \frac{e^3(x,t)}{12\eta}$, where $\eta$ is the solvent viscosity. Lateral flux $j_v$ is given by the imbalances between osmotic and hydraulic pressures with the exterior, such that

$$j_v = \lambda_v(\mathcal{R}T\delta c - \delta p),$$

with $\lambda_v$ the solvent permeability. Neglecting spatial variations in bridge thickness $e(x,t) \sim e_0$ (thus $\kappa_v = \kappa_v(e_0)$) yields the bridge mass balance Eq.3 in the main text:

$$\kappa_v \frac{\partial^2 \delta p}{\partial x^2} + 2\lambda_v \left[\mathcal{R}T\delta c(x,t) - \delta p(x,t)\right] = 0, \tag{7}$$

**Solute balance** The local solute balance in the bridge is given by

$$\begin{aligned}[\delta c(x,t+dt)e(x,t+dt) - \delta c(x,t)e(x,t)]\, dx = {} & [j_d(x,t)e(x,t) - j_d(x+dx,t)e(x+dx,t)]\, dt \\ & + 2j_s dt dx \left[1 + \left(\frac{\partial e}{\partial x}\right)^2\right]\end{aligned}$$

with the longitudinal diffusive flux given by Fick's law

$$e(x,t)j_d(x,t) = -e(x,t)\, D\frac{\partial \delta c}{\partial x},$$

$j_s$ is the lateral solute incoming flux per unit bridge length, composed of passive (p) and active (a) contributions

$$j_s = j^p + j^a = \mathcal{R}T\lambda_s \log\left(\frac{c_0}{c(x,t)}\right) + j^a,$$

with $\lambda_s$ the solute permeability coefficient

In the next, we assume that $\frac{\partial \delta c}{\partial t} \sim 0$, that is that the typical solute diffusion time $\tau_D = \ell_0^2/D$ is very small compared to all other timescales in the system (i.e. $\tau_{v,s}$). Neglecting spatial variations in bridge thickness $e(x,t) \sim e_0$ and assuming small variations in solute concentration $\delta c \ll c_0$, as above, yields the solute balance Eq.4 in the main text:

$$e_0 D \frac{\partial^2 \delta c}{\partial x^2} = 2\lambda_s \mathcal{R}T \frac{\delta c}{c_0} - 2j^a, \tag{8}$$

This equation corresponds to the continuity equation $e(x,t)\frac{D}{Dt}c(x,t) = -\partial_x J$, with flux $J = -D\partial_x c$ and the material derivative $\frac{D}{Dt} = \partial_t + v_x \partial_x$. Evaluation of the Peclet number for the flow generated by pressure difference between two lumens with a size difference of $L = 10\mu m$, separated by a distance $\ell = 1\mu m$ gives

$$\mathcal{P}e \equiv \frac{e_0.v_x}{D} \simeq 10^{-3} \tag{9}$$

showing that the advective part of the material derivative can be neglected with respect to diffusion/mass transport.

We give here an estimate for the active pumping flux. From [1], one has the estimated ion flux generated by the Na/K-ATPase active pumps to be $J_a \simeq 7.8 \times 10^6 ions/min/cell$, at the blastocyst stage, when the cavity is already formed. Assuming their activity is the same at the formation of the micro-lumens, we want to convert $J_a$ into the active pumping constant $j^a$, expressed as a flux of ions per unit time per unit area ($mol.s^{-1}.m^{-2}$).

At blastocyst stage, the blastocyst volume is estimated to be $0.66 \times 10^6 \mu m^3$ [1] and has almost spherical shape, giving an estimated surface $3.67 \times 10^4 \mu m^2$, for an average of 42 trophectoderm cells at this stage. This gives a surface of roughly $872\mu m^2$ per trophectoderm cell. Therefore, one obtains the estimation for the active pumping constant

$$j^a = \frac{7.8 \times 10^6 \text{ions}/(6.022 \times 10^{23} \text{mol}^{-1})}{(60s)(872 \times 10^{-12} m^2)} = 2.47 \times 10^{-10} mol.s^{-1}.m^{-2}$$

**Non-dimensionalization** Using the following dimensionless variables: $\delta p = \bar{p}\Pi_0$, $\delta c = \bar{c}c_0$, $x = x'\ell(t)$, $j^a = \bar{j}^a \lambda_s \mathcal{R}T$ yields the two dynamical equations for the bridge

$$\frac{\xi_s^2}{\ell(t)^2}\frac{\partial^2 \bar{c}}{\partial x'^2} = \bar{c}(x') - \bar{j}^a, \tag{10}$$

$$\frac{\xi_v^2}{\ell(t)^2}\frac{\partial^2 \bar{p}}{\partial x'^2} = \bar{p}(x') - \bar{c}(x'), \tag{11}$$

where we have introduced the dimensionless solute (s) and solvent (v) screening numbers $\chi_{s,v}^2(t) = \xi_{s,v}^2/\ell^2(t)$.

#### 1.3 Calculation of the boundary fluxes $J^{s,v}$

With (10) and (11), one can find the analytic expressions of the concentration and pressure profiles along the bridge, and therefore explicit formula for solute and solvent fluxes $J_i^{s,v}$ from a lumen.

**Concentration profile** The concentration equation (10) in the bridge is similar to a steady-state heat equation with a source term ($\bar{j}^a$) and a radiative term ($\bar{c}(x', t)$). A particular solution is $\bar{c}(x) = \bar{j}^a$ and boundary conditions are given by:

$$\bar{c}(x' = -\frac{1}{2}) = \delta\bar{C}_1 = \mu_1 \frac{\bar{N}_1}{\bar{L}_1^2} - 1, \tag{12}$$

$$\bar{c}(x' = \frac{1}{2}) = \delta\bar{C}_2 = \mu_2 \frac{\bar{N}_2}{\bar{L}_2^2} - 1, \tag{13}$$

which yields the following solution for $\bar{c}(x')$

$$\boxed{\bar{c}(x', t) = \bar{j}^a - (\delta\bar{C}_1 - \bar{j}^a)\frac{\sinh\left[\frac{x'-\frac{1}{2}}{\chi_s(t)}\right]}{\sinh(1/\chi_s(t))} + (\delta\bar{C}_2 - \bar{j}^a)\frac{\sinh\left[\frac{x'+\frac{1}{2}}{\chi_s(t)}\right]}{\sinh(1/\chi_s(t))},} \tag{14}$$

**Pressure profile** The pressure equation in the bridge (11) depends on the concentration profile $\bar{c}(x')$ above. Boundary conditions are given by

$$\bar{p}(x' = -\frac{1}{2}) = \delta\bar{P}_1 = \frac{\epsilon_1}{\bar{L}_1}, \tag{15}$$

$$\bar{p}(x' = \frac{1}{2}) = \delta\bar{P}_2 = \frac{\epsilon_2}{\bar{L}_2}, \tag{16}$$

The homogeneous solution is straightforward:

$$\bar{p}_H(x') = \lambda_0 e^{x'/\chi_v} + \mu_0 e^{-x'/\chi_v},$$

To find the particular solution, we use the parameter variation method, introducing the functions $\lambda(x')$ and $\mu(x')$ as

$$\bar{p}(x') = \lambda(x') e^{x'/\chi_v} + \mu(x') e^{-x'/\chi_v},$$

The expression of these functions are

$$\begin{cases} \lambda(x') & = \lambda_a + \int_a^{x'} dy E^-(y), \\ \mu(x') & = \mu_b + \int_b^{x'} dy E^+(y), \\ E^\pm(x') & = \pm\frac{\bar{c}(x')}{2\chi_v} e^{\pm x'/\chi_v}, \end{cases}$$

Using boundary conditions, and choosing $a = \frac{1}{2}, b = -\frac{1}{2}$, we get the final expression of $\bar{p}(x')$

$$\boxed{\begin{aligned} \bar{p}(x) = &-\frac{\delta\bar{P}_1 \sinh\left[\frac{x'-\frac{1}{2}}{\chi_v}\right]}{\sinh(1/\chi_v)} + \frac{\delta\bar{P}_2 \sinh\left[\frac{x'+\frac{1}{2}}{\chi_v}\right]}{\sinh(1/\chi_v)} + \lambda(x') e^{x'/\chi_v} + \mu(x') e^{-x'/\chi_v} + \\ &+ \frac{e^{-1/2\chi_v}}{\sinh 1/\chi_v}\left[\lambda(-\frac{1}{2})\sinh\left(\frac{x'-\frac{1}{2}}{\chi_v}\right) - \mu(\frac{1}{2})\sinh\left(\frac{x'+\frac{1}{2}}{\chi_v}\right)\right], \end{aligned}} \tag{17}$$

The expressions for $\lambda(x')$, $\mu(x')$ can be found analytically

$$\lambda(x') = \int_{\frac{1}{2}}^{x'} E^-(y)dy = \int_{\frac{1}{2}}^{x'} \left(\frac{-\bar{c}(y)}{2\chi_v} e^{-y/\chi_v}\right) dy = \frac{\bar{j}^a}{2}\left[e^{-x'/\chi_v} - e^{-1/2\chi_v}\right] + \frac{\delta\bar{C}_1 - \bar{j}^a}{2\chi_v s_1} I_1^-(x') - \frac{\delta\bar{C}_2 - \bar{j}^a}{2\chi_v s_1} I_2^-(x'),$$

$$\mu(x') = \int_{-\frac{1}{2}}^{x'} E^+(y)dy = \int_{-1/2}^{x'} \left(\frac{\bar{c}(y)}{2\chi_v} e^{y/\chi_v}\right) dy = \frac{\bar{j}^a}{2}\left[e^{x'/\chi_v} - e^{-1/2\chi_v}\right] - \frac{\delta\bar{C}_1 - \bar{j}^a}{2\chi_v s_1} I_1^+(x') + \frac{\delta\bar{C}_2 - \bar{j}^a}{2\chi_v s_1} I_2^+(x'),$$

where, defining

$$s_1 \equiv \sinh\frac{1}{\chi_s}; c_1 \equiv \cosh\frac{1}{\chi_s}; \qquad s^{\pm} \equiv \sinh\left(\frac{x' \pm \frac{1}{2}}{\chi_s}\right); c^{\pm} \equiv \cosh\left(\frac{x' \pm \frac{1}{2}}{\chi_s}\right)$$

we find

$$I_1^-(x) = \int_{1/2}^{x} dy e^{-y/\chi_v} \sinh\frac{y-\frac{1}{2}}{\chi_s} = \frac{\chi_v\chi_s}{(\chi_v-\chi_s)(\chi_v+\chi_s)}\left[e^{-x/\chi_v}\left(\chi_v c^-(x) + \chi_s s^-(x)\right) - \chi_v e^{-1/2\chi_v}\right], \tag{18a}$$

$$I_2^-(x) = \int_{1/2}^{x} dy e^{-y/\chi_v} \sinh\frac{y+\frac{1}{2}}{\chi_s} = \frac{\chi_v\chi_s}{(\chi_v-\chi_s)(\chi_v+\chi_s)}\left[e^{-x/\chi_v}\left(\chi_v c^+(x) + \chi_s s^+(x)\right) - e^{-1/2\chi_v}\left(\chi_v c_1 + \chi_s s_1\right)\right], \tag{18b}$$

$$I_1^+(x) = \int_{-1/2}^{x} dy e^{y/\chi_v} \sinh\frac{y-\frac{1}{2}}{\chi_s} = \frac{\chi_v\chi_s}{(\chi_v-\chi_s)(\chi_v+\chi_s)}\left[e^{x/\chi_v}\left(\chi_v c^-(x) - \chi_s s^-(x)\right) - e^{-1/2\chi_v}\left(\chi_v c_1 + \chi_s s_1\right)\right], \tag{18c}$$

$$I_2^+(x) = \int_{-1/2}^{x} dy e^{y/\chi_v} \sinh\frac{y+\frac{1}{2}}{\chi_s} = \frac{\chi_v\chi_s}{(\chi_v-\chi_s)(\chi_v+\chi_s)}\left[e^{x/\chi_v}\left(\chi_v c^+(x) - \chi_s s^+(x)\right) - \chi_v e^{-1/2\chi_v}\right], \tag{18d}$$

The denominator is not defined for $\chi_v = \chi_s$, so we calculate the functions $I_{1,2}^{\pm}$ for $\chi_s = \chi_v = X$

$$I_1^-(x) = \frac{1}{2}\left[\left(x-\frac{1}{2}\right)e^{-1/2X} + \frac{X}{2}\left(e^{-2x/X} - e^{-1/X}\right)e^{1/2X}\right], \tag{19a}$$

$$I_2^-(x) = \frac{1}{2}\left[\left(x-\frac{1}{2}\right)e^{1/2X} + \frac{X}{2}\left(e^{-2x/X} - e^{-1/X}\right)e^{-1/2X}\right], \tag{19b}$$

$$I_1^+(x) = \frac{1}{2}\left[\frac{X}{2}\left(e^{2x/X} - e^{-1/X}\right)e^{-1/2X} - \left(x+\frac{1}{2}\right)e^{1/2X}\right], \tag{19c}$$

$$I_2^+(x) = \frac{1}{2}\left[\frac{X}{2}\left(e^{2x/X} - e^{-1/X}\right)e^{1/2X} - \left(x+\frac{1}{2}\right)e^{-1/2X}\right], \tag{19d}$$

##### 1.3.1 Fluxes

**Solute fluxes** The dimensionless solute boundary fluxes $\bar{J}_{1,2}^s$ are given by the concentration $\bar{c}(x)$ as

$$\begin{cases} \bar{J}_1^s(t) &= -\frac{\bar{\xi}_s^2}{\ell}\partial_{x'}\bar{c}(x')|_{x'=-\frac{1}{2}}, \\ \bar{J}_2^s(t) &= \frac{\bar{\xi}_s^2}{\ell}\partial_{x'}\bar{c}(x')|_{x'=\frac{1}{2}}, \end{cases} \tag{20}$$

thus

$$\bar{J}_1^s = \bar{\xi}_s\left[(\delta\bar{C}_1 - \bar{j}^a)\coth\left(\frac{1}{\chi_s}\right) - (\delta\bar{C}_2 - \bar{j}^a)\frac{1}{\sinh(1/\chi_s)}\right], \tag{21a}$$

$$\bar{J}_2^s = \bar{\xi}_s\left[(\delta\bar{C}_2 - \bar{j}^a)\coth\left(\frac{1}{\chi_s}\right) - (\delta\bar{C}_1 - \bar{j}^a)\frac{1}{\sinh(1/\chi_s)}\right], \tag{21b}$$

**Solvent fluxes** The dimensionless solvent boundary fluxes $\bar{J}_{1,2}^v$ are given by continuity of the pressure

$$\begin{cases} \bar{J}_1^v &= -\frac{\bar{\xi}_v^2}{\ell}\partial_{x'}\bar{p}(x')|_{x'=-\frac{1}{2}}, \\ \bar{J}_2^v &= \frac{\bar{\xi}_v^2}{\ell}\partial_{x'}\bar{p}(x')|_{x'=\frac{1}{2}} \end{cases} \tag{22}$$

so that

$$\bar{J}_1^v = \bar{\xi}_v\left[\coth\left(\frac{1}{\chi_v}\right)\left(\delta\bar{P}_1 - \lambda(-1/2)\,e^{-1/2\chi_v}\right) - \frac{1}{\sinh(1/\chi_v)}\left(\delta\bar{P}_2 - \mu(1/2)\,e^{-1/2\chi_v}\right) - \lambda(-1/2)\,e^{-1/2\chi_v}\right], \tag{23a}$$

$$\bar{J}_2^v = \bar{\xi}_v \left[\coth\left(\frac{1}{\chi_v}\right)\left(\delta\bar{P}_2 - \mu\,(1/2)\,e^{-1/2\chi_v}\right) - \frac{1}{\sinh(1/\chi_v)}\left(\delta\bar{P}_1 - \lambda\,(-1/2)\,e^{-1/2\chi_v}\right) - \mu\,(1/2)\,e^{-1/2\chi_v}\right], \tag{23b}$$

**Limit case** In the limits of vanishing or very large solute/solvent screening lengths compared to the bridge length, one obtains :

- For large screening length $\xi_{s,v} \gg \ell(t)$, one has

$$\bar{J}_1^s \sim \frac{\bar{\xi}_s^2}{\bar{\ell}}\left[\delta\bar{C}_1 - \delta\bar{C}_2\right] \simeq -\bar{J}_2^s \tag{24}$$

$$\bar{J}_1^v \sim \frac{\bar{\xi}_v^2}{\bar{\ell}}\left[\delta\bar{P}_1 - \delta\bar{P}_2\right] \simeq -\bar{J}_2^v \tag{25}$$

- For small screening lengths $\xi_{s,v} \ll \ell(t)$, one has

$$\bar{J}_i^s \sim \bar{\xi}_s\left(\delta\bar{C}_i - \bar{j}^a\right), \quad i = 1,2 \tag{26}$$

$$\bar{J}_i^v \sim \bar{\xi}_v \delta\bar{P}_i, \quad i = 1,2 \tag{27}$$

###### 1.3.2 Net fluxes

The net osmotic and solvent fluxes are defined, for the lumens $1,2$, as $\bar{J}_{2\to1}^s = \bar{J}_2^s - \bar{J}_1^s$ and $\bar{J}_{2\to1}^v = \bar{J}_2^v - \bar{J}_1^v$. Using their expressions we find

$$\bar{J}_{2\to1}^s = \bar{\xi}_s \Delta C \coth\frac{1}{2\chi_s}, \tag{28}$$

$$\bar{J}_{2\to1}^v = \bar{\xi}_v\left[\Delta P.\coth\frac{1}{2\chi_v} + \Delta C.\frac{1}{\sinh(\frac{1}{2\chi_v})}.\Lambda(\chi_s,\chi_v)\right], \tag{29}$$

where $\Delta C = \delta\bar{C}_2 - \delta\bar{C}_1$, $\Delta P = \delta\bar{P}_2 - \delta\bar{P}_1$ and

$$\Lambda(\chi_s,\chi_v) = \frac{1}{\sinh\frac{1}{\chi_s}}\frac{\chi_s}{\chi_v^2 - \chi_s^2}\left[\chi_s s_1 \cosh\frac{1}{2\chi_v} - \chi_v(1+c_1)\sinh\frac{1}{2\chi_v}\right] \tag{30}$$

In the special case where $\chi_s = \chi_v = \chi$, the function $\Lambda$ reduces to

$$\Lambda(\chi) = \frac{1 - \chi\sinh(\frac{1}{\chi})}{2\chi\sinh(\frac{1}{\chi})}\cosh\left(\frac{1}{2\chi}\right) \tag{31}$$

###### 1.3.3 Moving boundary fluxes

One shall take special care into dealing with the moving boundary problem (Stefan problem). In our case, the border of the lumen moves due to incoming flux, hence the need to state carefully the system. As represented on Fig. S2 we restrict ourselves here to half a lumen, since it is symmetric, and focus on the hydraulic limit. The lumen is centered at $X_0$, with length $L$ corresponding to the position $X_\ell$ of its right border. The bridge starts at $X_\ell(t)$ and stops at position $X_B$. Both $X_0, X_B$ are supposed fixed positions in the reference frame.

First we shall express the half lumen area ($A_{\text{Lumen}}/2$), assuming the lumen is symmetric. This can be expressed as

$$\frac{1}{2}A_{\text{Lumen}}(t) = A_{1/2}(t) + e_0(X_\ell(t) - X_0)$$

but here we do not assume that $e_0 \sim 0$. The bridge has and area $A_B = e_0(X_B - X_\ell(t)$ The lumen generates an outgoing flux $J(t)$, whil the bridge generates an outgoing flux $J_B(t)$. Assuming no variation in diameter ($e_0 = cte$) of the bridge, the fluxes are equal ($J(t) = J_B(t)$). The total mass/area conservation reads

$$\frac{1}{2}A_{\text{Lumen}} + A_B = -\int dt J_B(t) \tag{32}$$

Thus, derivation of the above equation wrt time leads to

$$\frac{dA_{1/2}}{dt} + e_0\frac{dX_\ell}{dt} - e_0\frac{dX_\ell}{dt} = -J_B$$

and finally

$$\frac{dA_{1/2}}{dt} = -J_B \tag{33}$$

The flux $J_B$ can be expressed as a function of boundary pressures, given it is a Poiseuille flow. Therefore, in spite of the moving character of the boundary, we recover exactly the expression expected for mass conservation in the simplest case of pure hydraulic exchanges without permeability nor osmotic effects. The same reasoning could be applied for this more complex situation as well as for boundary solute fluxes.

### 2 Chain of micro-lumens

We detail here the analytical results for a one-dimensional chain of connected micro-lumens.

#### 2.1 Hydraulic limits

We show here how one may reduce the hydro-osmotic model equations for length $L_i$ and number of solutes $N_i$ to a single equation in the appropriate limits.

##### 2.1.1 Without active pumping

We start with two microlumens, labeled $i, j$, connected by a bridge of dimensionless length $\bar{\ell}_{ij}$. We consider first the case where lumens do not pump solutes ($\bar{j}^a = 0$). Assuming (i) there is no pressure screening $\chi_v \gg 1$, (ii) that solute concentrations are screened $\chi_s \ll 1$ and (iii) that the relaxation time for the number of solutes within lumens is negligible compared to the water relaxation time, $\tau_s \ll \tau_v$, the dimensionless equations Eq. [6-7] reduce to

$$\frac{d\bar{L}_i}{dt} = \frac{\mu\nu}{\tau_v}\left[\delta\bar{C}_i - \frac{\epsilon}{\bar{L}_i}\right] - \frac{\mu}{2\tau_v\bar{L}_i}\frac{\bar{\xi}_v^2}{\bar{\ell}_{ij}}\left[\frac{\epsilon}{\bar{L}_i} - \frac{\epsilon}{\bar{L}_j}\right] \tag{34a}$$

$$\frac{\tau_s}{\tau_v}\frac{d\bar{N}_i}{dt} = -\frac{2\nu\bar{L}_i}{\tau_v}\delta\bar{C}_i \tag{34b}$$

where we used the expressions of fluxes (25) and (26). Because $\tau_s \ll \tau_v$, the left-hand side of the second equation is zero, thus $\delta\bar{C}_i = 0$, the lumens relax immediately to osmotic equilibrium. Since we have $\epsilon \ll 1$, the first term of the right-hand side of the first equation is negligible. Finally, we are left with

$$\frac{d\bar{L}_i}{dt} = -\frac{\mu\epsilon\bar{\xi}_v^2}{2\tau_v\bar{L}_i\bar{\ell}_{ij}}\left[\frac{1}{\bar{L}_i} - \frac{1}{\bar{L}_j}\right] = \frac{1}{T_h\bar{L}_i\bar{\ell}_{ij}}\left[\frac{1}{\bar{L}_j} - \frac{1}{\bar{L}_i}\right] \tag{35}$$

where we define the typical hydraulic time

$$T_h \equiv \frac{2\tau_v}{\mu\epsilon\bar{\xi}_v^2} = \frac{2\tau_v\ell_0 L_0}{\mu\epsilon\xi_v^2} \tag{36}$$

###### 2.1.2 With active pumping

Consider now the case where active pumping is non-zero $\bar{j}^a \neq 0$. Assuming the same limits as before yields now

$$\frac{d\bar{L}_i}{dt} = \frac{\mu\nu}{\tau_v}\delta\bar{C}_i - \frac{\mu}{2\tau_v\bar{L}_i}\frac{\bar{\xi}_v^2}{\bar{\ell}_{ij}}\left[\frac{\epsilon}{\bar{L}_i} - \frac{\epsilon}{\bar{L}_j}\right] \tag{37a}$$

$$0 = \frac{2\nu\bar{L}_i}{\tau_v}\left[\bar{j}^a - \delta\bar{C}_i\right] \tag{37b}$$

Injecting (37b) into (37a) the system of equations reduces to

$$\frac{d\bar{L}_i}{dt} = \frac{\mu\nu}{\tau_v}\bar{j}^a - \frac{\mu\epsilon\bar{\xi}_v^2}{2\tau_v\bar{L}_i\bar{\ell}_{ij}}\left[\frac{1}{\bar{L}_i} - \frac{1}{\bar{L}_j}\right] = \frac{1}{T_h\bar{L}_i\bar{\ell}_{ij}}\left[\frac{1}{\bar{L}_j} - \frac{1}{\bar{L}_i}\right] + \frac{1}{T_p} \tag{38}$$

where we have defined the typical pumping time

$$T_p \equiv \frac{\tau_v}{\mu\nu\bar{j}^a} \tag{39}$$

##### 2.2 Mean-field theory of lumen coarsening

###### 2.2.1 Derivation of the scaling law

We present here a mean-field derivation of the scaling law for the number of lumens $\mathcal{N}$ for a linear chain embedded in a space of dimension $d$. This was first done in [2] in the case $d = 2$, and we adapt it here to our model, with the lumen size denoted $X$ ($X = A$ in $d = 2$; $X = V$ in $d = 3$). We consider a chain of lumens of total length $\mathcal{L}_0$, assumed to be constant, without active pumping. Let $\phi(X,t)$ be the distribution of drops with size $X$ at time $t$, such that $\int_X^{X+\Delta X} dX'\phi(X',t)$ is the number of lumens with lsizes between $X$ and $X + \Delta X$. The total number of lumens at time $t$ is

$$\mathcal{N}(t) = \int_0^\infty dX\phi(X,t), \tag{40}$$

The total mass is defined as $M_{\text{tot}} = \sum_i X_i$ in the discrete case, corresponding to the first moment of the distribution. In continuous form, we have

$$M_{\text{tot}} = \int_0^\infty dXX\phi(X,t) \tag{41}$$

The total mass is analoguous to total area in $d = 2$, and to the total volume in $d = 3$. In the case of a purely hydraulic chain, the conservation law is written as

$$\partial_t\phi + \partial_X\left[\phi u\right] = 0, \tag{42}$$

with $u = \frac{dX}{dt}$ the rate of change in size of a lumens, given in $d$ dimension by

$$\frac{dX}{dt} = \frac{1}{\bar{\ell}}\left[X_*^{-1/d} - X^{-1/d}\right], \tag{43}$$

where $X_*$ is the mean-field size of the system, defined as $X_*^{-1/d} = \frac{1}{\mathcal{N}}\int_0^\infty dXX^{-1/d}\phi(X,t)$, the harmonic average of the lumen size. The typical distance between two lumens $\bar{\ell}$, or mean separation, can also be expressed as $\bar{\ell} = \mathcal{L}_0/\mathcal{N}(t)$, and, by definition, the average size of the lumens is defined as $\bar{X} = \frac{1}{\mathcal{N}}\int X\phi(X,t)dX = M_{\text{tot}}/\mathcal{N}(t)$.

Thus, $\bar{\ell} = \bar{X}\mathcal{L}_0/M_{\text{tot}}$, with the average size $\bar{X}$, which scales like $X_*$, such that $\gamma = X_*/\bar{X}$. $\gamma$, $\mathcal{L}_0$, $M_{\text{tot}}$ are supposed to be constant.

We assume that the chain is in the regime in which the distribution $\phi$ is self-similar, writing it as a scale-invariant self-similar distribution, $\phi(X,t) = t^{-\alpha}f_d(z)$, $z = \frac{X}{X_*}$ and $d$ is the dimension. The mean-field length also has a scaling form $X_* = \sigma t^\beta$, with undetermined constants $\sigma$, $\alpha$, $\beta$ and $\gamma$.

The total mass is written as

$$M_{\rm tot} = \sigma^2 t^{2\beta-\alpha} \int_0^{\infty} dz z f_d(z), \tag{44}$$

and the total mass conservation yields $2\beta = \alpha$. Expanding the conservation law terms leads to

$$\beta t^{-2\beta-1}\left[2f_d(z) + zf_d'(z)\right] = \frac{M_{\rm tot}\gamma}{\mathcal{L}_0\sigma^{2+1/d}} t^{-4\beta-\frac{\beta}{d}} \frac{d}{dz}\left[f_d(z)\left(1 - z^{-\frac{1}{d}}\right)\right] \tag{45}$$

$$\frac{g_1(z)}{g_2(z)} = \frac{C}{\beta} t^{-2\beta-\frac{\beta}{d}+1}, \tag{46}$$

where $g_1(z) = 2f_d(z) + zf_d'(z)$, $g_2(z) = \frac{d}{dz}\left(f_d(z)(1 - z^{-1/d})\right)$, $C = \frac{M_{\rm tot}\gamma}{\mathcal{L}_0\sigma^{2+1/d}}$. Since $z$ and $t$ are assumed to be independent variables, then we have $\beta = \left(2 + \frac{1}{d}\right)^{-1}$, and normalizing the function $f$, as $\int_0^{\infty} dz f(z) = 1$, we find the power-law for the number of lumens as

$$\boxed{\mathcal{N}(t) = \sigma t^{-\beta}, \beta = \frac{1}{2 + \frac{1}{d}}} \tag{47}$$

For $d = 2$, we find $\beta = \frac{2}{5}$ [2], and for $d = 3$, one has $\beta = \frac{3}{7}$, in agreement with [3] for a linear network of 3D-droplets. Similar arguments can be used to derive the scaling law $\mathcal{N}(t) \propto t^{-3/4}$ for 3D-droplets on a 2D network (instead of a linear chain) [3–5].

##### 2.2.2 Mean-field distribution of lumen size

From (47), one may express the function $f_d$ as relation between $f_d(z)$ and its derivative $f_d'(z)$ as :

$$\frac{df_d}{dz} = \left[\frac{\frac{4C}{d\beta} - 8z^{\frac{d+1}{d}}}{4z^{\frac{2d+1}{d}} + \frac{4C}{\beta} z\left(1 - z^{1/d}\right)}\right] f_d(z) \tag{48}$$

In the case $d = 2$, this reduces to the similarity ODE found by [2], and an explicit solution can be found in [6]. This expression is valid in for $z \in [0, z_{\rm max}[$, and one may find expressions of $C$ and $z_{\rm max}$ assuming $z_{\rm max}$ is an irregular singular point, such that the denominator and its derivative vanish for $z = z_{\rm max}$. We find the general expressions to be

$$z_{\rm max} = \left(\frac{d+1}{d}\right)^d; \quad C = \beta d \left(\frac{d+1}{d}\right)^{d+1} \tag{49}$$

which gives for $d = 2$, $\beta = 2/5$, $z_{\rm max} = \frac{9}{4}$ and $C = \frac{27}{10}$. The function $f_2$ is plotted on Fig. 4C.

##### 2.2.3 Coalescence scaling law

We propose the argument for derivation of the coalescence scaling law for the number of lumens $\mathcal{N}(t) \sim t^{-1}$ in any dimension $d$. We consider two droplets $i, j$ of radius $R_i$ and total mass $X_i \propto R_i^d$, at a distance $\ell_{ij}$. The total mass is analoguous to the area for $d = 2$ ($X_i = A_i \propto R_i^2$) and to the volume for $d = 3$ ($X_i = V_i \propto R_i^3$). Assuming that the distribution of active pumps is uniform over the surface of exchange $\mathcal{S}_i \propto R_i^{d-1}$ (the lumen perimeter for $d = 2$ and the lumen surface for $d = 3$), then the mass conservation is written as

$$\frac{dX_i}{dt} = j^a \mathcal{S}_i \Leftrightarrow dR_i^{d-1}\frac{dR_i}{dt} = j^a R_i^{d-1} \Leftrightarrow \frac{dR_i}{dt} = j^a$$

The radius increases linearly with $j^a$, $R_i \sim j^a.t$ and the distance between droplets $i, j$ decreases linearly in time, $\ell_{ij} \sim -j^a.t$ as the lumens do not move, and they coalesce when $\ell_{i,j} = 0$, hence the number of lumens decreases at a rate

$$\mathcal{N}(t) \sim (t/T_p)^{-1} \tag{50}$$

where $T_p$ is the typical pumping time given in (39). Therefore, the scaling exponent for coalesence dominated regime is independent from the dimension of the system.

##### 2.2.4 Size distribution of microlumens on the active pumping plateau

We consider a chain with uniform active pumping in the hydraulic limit, and we assume that we are in the regime $t > T_p$. In this regime, we study the part where the number of lumens remains roughly constant, $\mathcal{N}(t) \sim \mathcal{N}_p$, before the coalescence dominated regime. The evolution equation for a single lumen $i$ can be written as $\frac{dL_i}{dt} = \frac{1}{T_p}$. The solution is immediately found to be $L_i(t) = L_i(0) + \frac{t}{T_p}$, where time $t = 0$ is the time at which we consider the start of the regime. The population of lumens corresponds to a distribution with time-dependent mean $\mu(t)$ and standard deviation $\sigma(t)$. By definition, at a given time, one has

$$\mu(t) = \frac{1}{N_p} \sum_{i=1}^{N_p} L_i(t) = \mu(0) + \frac{t}{T_p} \tag{51a}$$

$$\sigma^2(t) = \frac{1}{N_p} \sum_{i=1}^{N_p} (L_i(t) - \mu(t))^2 = \sigma^2(0) \tag{51b}$$

The size distribution of lumens can be approximated by a Gaussian distribution (most likely distribution) with mean $\mu(t)$ and standard deviation $\sigma(t)$ :

$$\phi(L,t) \sim \exp\left[-\frac{(L(t) - \mu(t))^2}{\sigma^2(t)}\right] = \exp\left[-\frac{(x(t) - 1)^2}{a^2(t)}\right] \tag{52}$$

where $a^2(t) = \mu^2(t)\sigma^2(t)$ and $x(t) = \frac{L(t)}{\mu(t)}$. Now, in the limit $t \gg 1$, but such that no coalescence nor coarsening event happens,

$$\lim_{t\to\infty} x(t) = 1; \lim_{t\to\infty} a(t) = \infty$$

and the distribution tends to a Dirac distribution :

$$\phi(L,t) \to \delta\left(\frac{L(t)}{\mu(t)} - 1\right)$$

This shows that on the plateau following hydraulic coarsening and coalescence-dominated regime, the distribution evolves to a Dirac distribution centered on the rescaled length $L/\bar{L}$.

##### 2.2.5 Heterogeneous pumping profile

We study the case of a composed pumping profile : a uniform pumping plus a localized gaussian perturbation. Let be $\bar{j}^a(x)$ the pumping profile, defined as

$$\forall x \in [0,1], \bar{j}^a(x) = \bar{j}^a_0 + \delta\bar{j}^a e^{-\frac{(x-\mu)^2}{\sigma^2}} \tag{53}$$

where the chain has been rescaled by its total length $L_{\text{tot}}$. The total pumping along the chain, $I_{\text{tot}}$, is given integrating the function along the chain, as

$$I_{\text{tot}} = \int_{x_{\min}}^{x_{\max}} \bar{j}^a(x)dx \tag{54}$$

since in practice, lumens can sometimes not be distributed along the full chain but only within $x_{\min}$ and $x_{\max}$. The effect of localized active pumping is measured comparing the total pumping of two regions : the perturbed (P) versus the constant (C) profile. This is done by the separation of the total pumping into two contributions, provided that we define the perturbed region to lies within $[\mu - n\sigma, \mu + n\sigma]$, where $n$ is an arbitrary integer.

$$I_{\text{tot}} = I_C + I_P = \left[\int_{x_{\min}}^{\mu-n\sigma} \bar{j}^a_0 dx + \int_{\mu+n\sigma}^{x_{\min}} \bar{j}^a_0 dx\right] + \int_{\mu-n\sigma}^{\mu+n\sigma} \left(\bar{j}^a_0 + \delta\bar{j}^a e^{-\frac{(x-\mu)^2}{\sigma^2}}\right) dx \tag{55}$$

with

$$I_C = \int_{x_{\min}}^{\mu-n\sigma} \bar{j}_0^a dx + \int_{\mu+n\sigma}^{x_{\max}} \bar{j}_0^a dx = \bar{j}_0^a \left[\Delta x - 2n\sigma\right] \tag{56}$$

and

$$I_P = \int_{\mu-n\sigma}^{\mu+n\sigma} \left( \bar{j}_0^a + \delta\bar{j}^a e^{-\frac{(x-\mu)^2}{\sigma^2}} \right) dx = 2n\sigma\bar{j}_0^a + \delta\bar{j}^a \sigma\sqrt{\pi}\mathrm{erf}(n) \tag{57}$$

where $\Delta x = x_{\max} - x_{\min}$, and $\mathrm{erf}(x)$ is the error function. Thus, the perturbed region has larger pumping than the constant region if

$$\begin{aligned} I_P > I_C &\Leftrightarrow 2n\sigma\bar{j}_0^a + \delta\bar{j}^a\sigma\sqrt{\pi}\mathrm{erf}(n) > \bar{j}_0^a \left[\Delta x - 2n\sigma\right] \\ &\Leftrightarrow \delta\bar{j}^a\sigma\sqrt{\pi}\mathrm{erf}(n) > \bar{j}_0^a \left[\Delta x - 4n\sigma\right] \\ &\Leftrightarrow \frac{\delta\bar{j}^a\sigma\sqrt{\pi}\mathrm{erf}(n)}{\Delta x - 4n\sigma} > \bar{j}_0^a \end{aligned}$$

which gives us the threshold $\bar{j}*$ as

$$\bar{j}_* = \frac{\delta\bar{j}^a\sigma\sqrt{\pi}}{(x_{\max} - x_{\min}) - 4n\sigma}\mathrm{erf(n)} \tag{58}$$

In Fig. 6, we have $n = 2$, $x_{\min} = 0.2, x_{\max} = 0.8$, $\sigma = 0.05$, $\delta\bar{j}^a = 1$, that gives $\bar{j}^* \simeq 0.44$.

### 3 Chain integration

#### 3.1 Chain generation

Chains are generated with normal distributions for the areas of lumens, $A_i \hookrightarrow \mathcal{N}(A_0, \sigma_A)$ and bridges $\ell_{i,j} \hookrightarrow \mathcal{N}(\ell_0, \sigma_b)$, with usually $A_0 = 1$ ($L_0 = \sqrt{\mu A_0} \simeq 0.78$), $\ell_0 = 10$ (indicated otherwise). The total length of the chain is given by $\mathcal{L}_0 = 2\sum_i L_i + \sum_{i,j} \ell_{i,j}$ and its conservation is used as a check for topology. The number of ions in a lumen is denoted $N_i$, and is choosen at osmotic equilibrium such that $N_i = \frac{\mathcal{L}_i^2}{\mu_i}$ or randomly with normal distribution $\mathcal{N}(N_0, \sigma_N)$.

The screening numbers $\chi_{s,v}$ are considered as input parameters, and the non-dimensionalized screening lengths $\xi_{s,v}$ are calculated according to the average bridge length $\ell_0$.

##### 3.1.1 Numerical scheme

The algorithm is composed of two important parts : the integration of ordinary differential equations (ODE) that solves the dynamic of the chain, and the topology, that handles events when a cavity disappear or fuse with another. It is schematically represented in Fig. S5.

**Initialization** First, the chain is initialized with the imported parameters from a config file. This is done using a homemade parser. Once the parameters imported, the chain is generated either randomly (with a seed if specified) or according to the given files (see examples in _configfiles folder). The status of the chain, either hydraulic (H) or hydroosmotic (HO), must be specified. Pumping profile (if specified) is also computed at that time.

**Integration** After generation, the ODEs are solved iteratively, with a given solver, either Runge-Kutta 45 (RK45) or Runge-Kutta-Felhberg 45 (RKF45, default).

**The RKF45 method** Numerical integration for a system of ODE with initial value problem (IVP) often uses Runge-Kutta (RK) methods, based of iterative integration. The classical method is known as RK45. In order to improve integration, Fehlberg [7] proposed a new way to obtain a fourth-order method, with error of order 5, hence known as Runge-Kutta-Felhberg or RKF45. If we consider the IVP

$$\frac{dy}{dt} = f(t, y(t)), \quad y(t=0) = y_0$$

with $y(t)$ a time-dependent variable and $f$ is a given function. The time interval may be discretized with a time-step $\Delta t$, such that $t_{i+1} = t_i + \Delta t$, and similarly $y_i = y(t_i)$. RKF45 proposes a first estimation for the next value $y_{i+1}$ given the time step and previous step $y_i$, as

$$y_{i+1} = y_i + \frac{25}{216}k_1 + \frac{1408}{2565}k_3 + \frac{2197}{4104}k_4 - \frac{1}{5}k_5,$$

with the weights

$$\begin{cases} k_1 & = \Delta t f(t, y_i) \\ k_2 & = \Delta t f(t_i + \frac{1}{4}\Delta t, y_i + \frac{1}{4}k_1) \\ k_3 & = \Delta t f(t_i + \frac{3}{8}\Delta t, y_i + \frac{3}{32}k_1 + \frac{9}{32}k_2) \\ k_4 & = \Delta t f(t_i + \frac{12}{13}\Delta t, y_i + \frac{1932}{2197}k_1 - \frac{7200}{2197}k_2 + \frac{7296}{2196}k_3) \\ k_5 & = \Delta t f(t_i + \Delta t, y_i + \frac{439}{216}k_1 - 8k_2 \frac{3680}{513}k_3 - \frac{845}{4104}k_4) \\ k_6 & = \Delta t f(t_i + \frac{1}{2}\Delta t, y_i - \frac{8}{27}k_1 + 2k_2 - \frac{3544}{2565}k_3 + \frac{1859}{4104}k_4 - \frac{11}{40}k_5) \end{cases}$$

**Error evaluation and Adaptive time-stepping** RKF45 method also finds another estimate of the next step $y_{i+1}$, that we will denote $z_{i+1}$, and is given by

$$z_{i+1} = y_i + \frac{16}{135}k_1 + \frac{6656}{12825}k_3 + \frac{28561}{56430}k_4 - \frac{9}{50}k_5 + \frac{2}{55}k_6,$$

Therefore, we can define an error $\epsilon$ as

$$\epsilon = |y_i - z_i|,$$

For large variations of the variable $y(t)$, the two estimates give a large error, and conversly, a small error for small variations. This error allows us to calculate a new time step $\Delta t_{i+1}$ such that [8]

$$\Delta t_{i+1} = \begin{cases} \Delta t_i S \left(\frac{\tau}{\epsilon}\right)^{0.2}, & \text{if } \epsilon > \tau \\ \Delta t_i S \left(\frac{\tau}{\epsilon}\right)^{0.25}, & \text{if } \epsilon \leq \tau \end{cases} \tag{59}$$

where $\Delta t_i$ is the previous time-step, $S$ is a security factor (arbitrary factor, usually $S = 0.9$), $\tau$ is the tolerance (arbitrary factor, usually $\tau \sim 10^{-6}$ in the litterature, we use $\tau \sim 10^{-10}$ to prevent divergences). Thus, if the error is higher than the tolerance, the time-step is reduced, otherwise it increases.

For both $RK45$ and $RKF45$ cases, we have one set of weights ($\{k^{(j)}\}$ for $L_j$'s) for H-chain, or two sets ($\{k^{(j)}, q^{(j)}\}$ for $L_j$'s and $N_j$'s) for HO-chain. Note that because the lumens are not moving during integration, the constraint $L_a + L_b + \ell_{ab} = \text{cste}$ must be enforced in the calculation of the weights. Since topological events modify the ODE equations, one cannot let them happen during integration. At the end of the weights calculation, a check is performed on the lengths of bridges and lumens. While topological event happens (merge or collapse), the integration is restarted from the previous time-step with a new halved time-step. This is a costly procedure, but required in order to avoid wrong results. The new time step is calculated at the end of the integration. For RK45, it is given by $\Delta t_{i+1} = \alpha / N(t_i^\beta)$. For RKF45, it is given according to (59). The error $\epsilon$ is calculated over the estimations of the lengths $L_j$, since divergences occur for at small lengths due to concentration and pressure terms. For estimations $y_{i+1}(L_j)$ and $z_{i+1}(L_j)$ of the length $L_j$ at step $i+1$, the error $\epsilon$ is the maximum of the $\epsilon_j$'s :

$$\epsilon = \max_j \epsilon_j = \max_j |y_{i+1}(L_j) - z_{i+1}(L_j)|, \tag{60}$$

$\tau$ can be changed in the config file. The security factor $S = 0.90$ is a constant.

**Topology** When an integration step is complete and valid, topology is checked. We assume all topological events to be instantaneous.

1. We first check whether lumens disappear. A lumen $i$ collapses if its length is below a given threshold $L_{\text{dis}}$
$$L_i \leq L_{\text{dis}}, \tag{61}$$
The value of $L_{\text{dis}}$ is defined in the config file. We choose $L_{\text{dis}} = 0.1$ in order to avoid divergence of the concentration when a lumen shrinks. When a lumen collapses, it is removed from the graph, as well as the bridges that connected it. Its two neighbors are connected via a new bridge, but they do not move.

2. We then check whether a lumen collides with the chain borders. If a lumen $i$ overlaps the border, it will be pushed towards the center, say $\ell_{0i}$ at left border. This is done by checking the length of the bridge between the lumen $i$ and the border, considered as a lumen. Assuming the lumen $i$ touches the left border (index 0), the new position $x'_i$ of the center of mass of the lumen is moved towards the center if
$$\ell_{0i} < 0 \Rightarrow x'_i = x_i + \ell_{0i}, \tag{62}$$
This works the same for the right border (index $-1$), such that $\ell_{i,-1} < 0 \Rightarrow x'_i = x_i - \ell_{i,-1}$.

3. Finally, we detect collisions between lumens. Two colliding lumens merge together, forming a new lumen. A collision between parent lumens $i$, $j$ is detected when
$$\ell_{ij} \leq \ell_{\text{merge}}, \tag{63}$$
where $\ell_{\text{merge}}$ is a length that is defined in the config file. We usually choose $\ell_{\text{merge}} = 10^{-3}$. When a collision occurs, a new lumen is created with index $k$, while parent lumens $i$, $j$ are deleted from the graph. The center of mass of the new lumen $k$ is given by the barycenter of the two parent lumens:
$$x_k = \frac{x_i A_i + x_j A_j}{A_i + A_j}, \tag{64}$$
where $x$ is the position and $A$ the area (equivalent to mass) of a lumen. The area of new lumen $k$ is calculated to conserve mass:
$$A_k = A_i + A_j, \tag{65}$$
and its length is calculated accordingly. If the lumen is hydro-osmotic, its total number of ions is conserved:
$$N_k = N_i + N_j, \tag{66}$$
Finally, if a profile $f_p(x)$ of active pumping is imposed, the new active pumping $j^a_k$ is given by the position $x_k$, such that
$$j^a_k = \frac{1}{|x_2 - x_1|} \int_{x_1}^{x_2} f_p(x) dx,$$
where $x_1, x_2$ are the positions of the borders of the lumen or the bridge. Otherwise, it is the average of the active pumping constants of merging lumens, $j^a_k = \frac{1}{2}(j^a_i + j^a_j)$. New bridges between neighbors of lumens $i$ and $j$ are created and their lengths are given by $L_k$ and $x_k$.

4. Since multiple lumens could merge at the same time, a loop detects whether there are still collisions occurring after the merging of lumens and redefinition of the connectivity graph.

At the end of the topology check, the algorithm checks whether stop conditions are fulfilled. Usually, the integration is stopped when $\mathcal{N}(t) \leq 1$, or if one lumen occupies the whole chain.
