## Supporting Figures for "A hydro-osmotic coarsening theory of biological cavity formation"

### Supporting S1 to S5 Figures

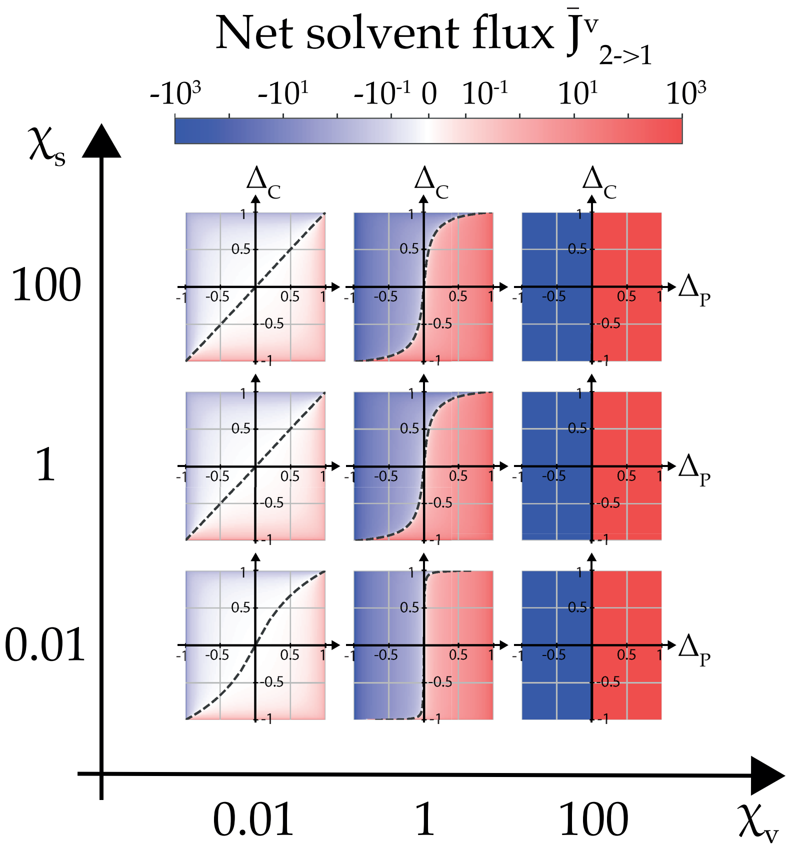


**S1 Fig. Net solvent flux diagrams.** Net solvent flux diagrams as function of the relative concentration asymmetry $\Delta_C$ and pressure asymmetry $\Delta_P$ for a 2-lumen system, at different values of the rescaled screening lengths $\chi_{s,v}$. Dashed lines represent the zero net flux $\bar{J}^v_{2\to1} = 0$. At large pressure screening length $\chi_v = 100$, the net flux does not depends on the concentration screening length $\chi_s$. For smaller pressure screening lengths $\chi_v = 0.01$, 1, we observe a mild influence of the concentration screening length when it approaches low values $\chi_s = 0.01$. This shows that the concentration screening length has limited influence on the net solvent fluxes compared to the pressure screening length. Asymmetry ratios are defined in the main text.

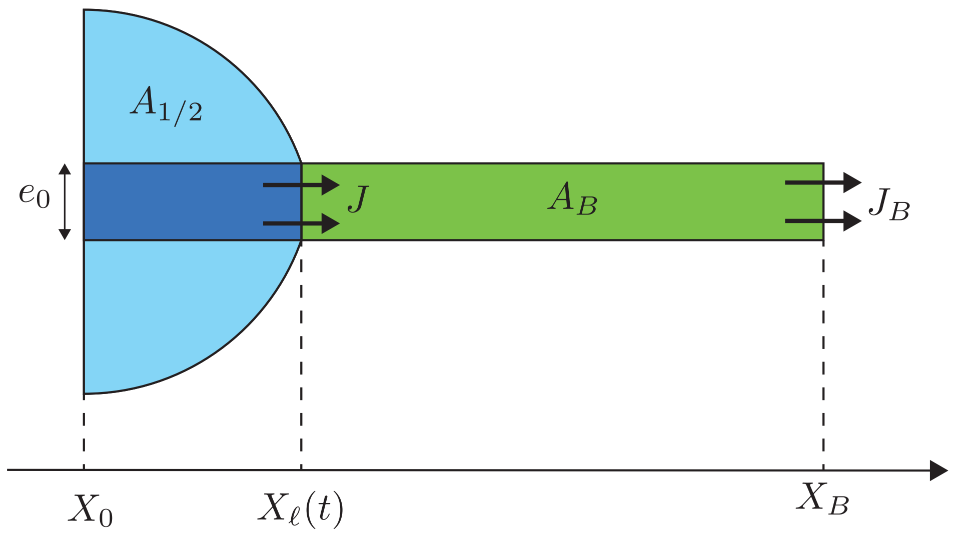


**S2 Fig. Half a lumen with moving boundary $X_\ell$.** The area of half the lumen is $A_{1/2}$ (light blue region) plus the bridge-lumen portion (dark blue region). The green region corresponds to the bridge area.

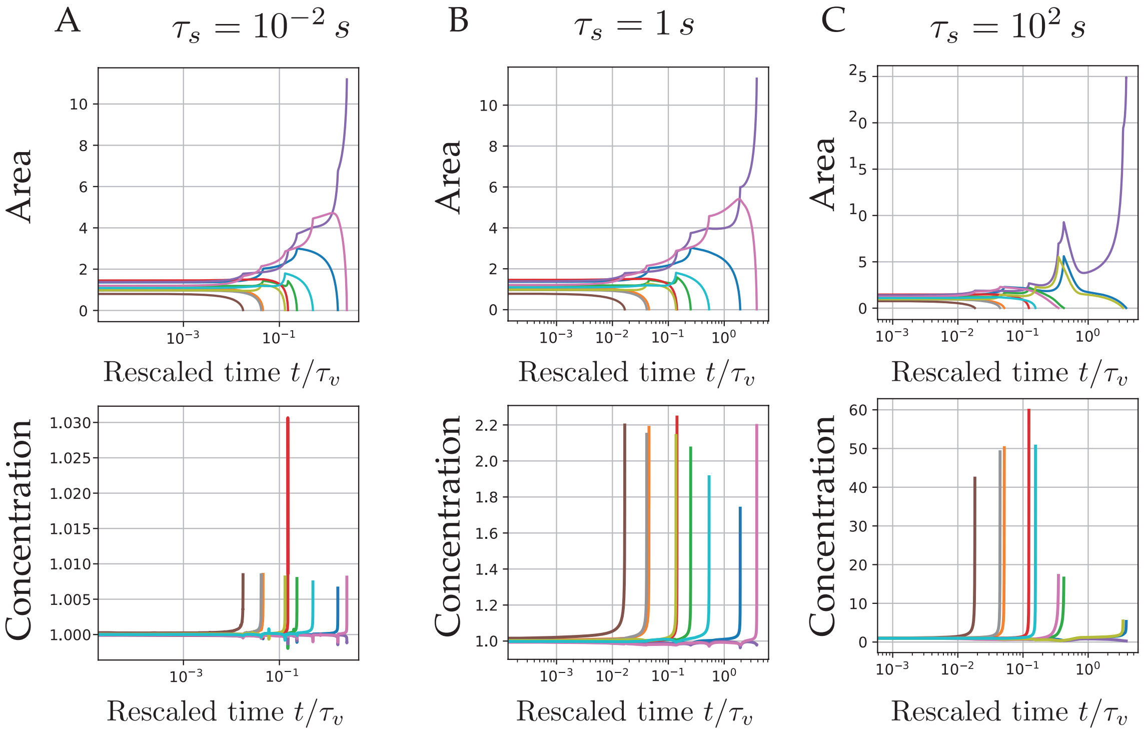


**S3 Fig. Typical dynamics of a chain with $\mathcal{N}(0) = 10$ lumens.** (A) With parameters $A_0 = 1$, $\ell_0 = 10$, $\chi_v = 50$, $\chi_s = 5$, $\tau_v = 1\, s$, $\tau_s = 0.01\, s$. (B) With parameters $\chi_v = 50$, $\chi_s = 5$, $\tau_v = 1\, s$, $\tau_s = 1\, s$. (C) With parameters $\chi_v = 50$, $\chi_s = 5$, $\tau_v = 1\, s$, $\tau_s = 100\, s$.

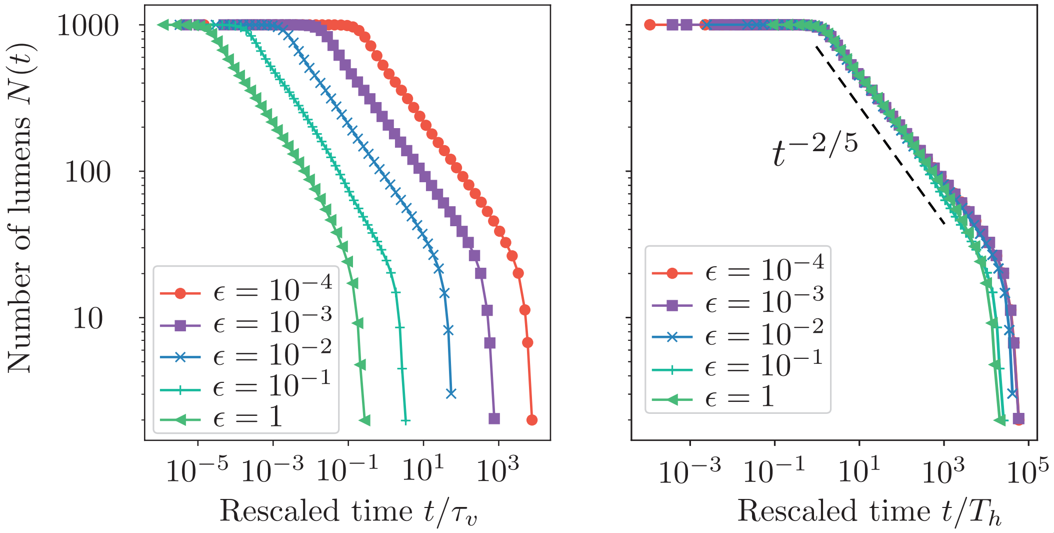


**S4 Fig. Influence of cell mechanics on coarsening.** The parameter $\epsilon$ is varied from $10^{-4}$ to 1. (Left) Number of lumens as function of the rescaled time $t/\tau_v$, showing the influence of $\epsilon$ on the hydraulic time $T_h$. (Right) Number of lumens as function of the rescaled time $t/T_h$. All the curve collapse but a small deviation from the scaling law $t^{-2/5}$ is observed for $\epsilon \gtrsim 10^{-1}$. The scaling law $\mathcal{N}(t) \sim t^{-2/5}$ is shown as a dashline. Parameters are $\chi_v = 50, \chi_s = 5, \tau_s = \tau_v = 1s$ and $\mathcal{N}(0) = 1000$. Each curve is an average of 20 simulations.

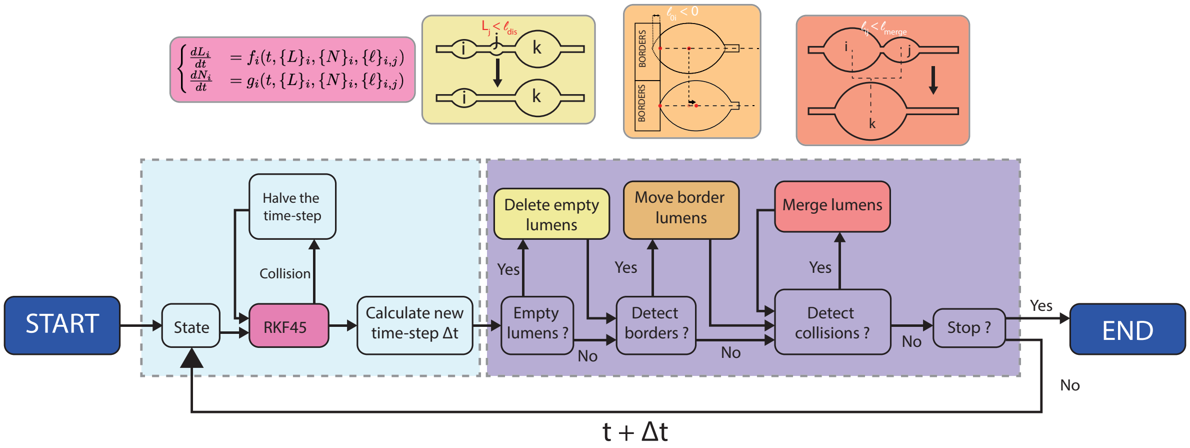


**S5 Fig. Representation of the numerical scheme.**
