## Supporting Table 1 for "A hydro-osmotic coarsening theory of biological cavity formation"

| **Name** | **Symbol** | **Units [2D]** | **Range [3D]** | **Units [3D]** |
|---|---|---|---|---|
| Thermal energy per unit mole | $RT$ | $N.m.mol^{-1}$ | $2.4 \times 10^3$ | $N.m.mol^{-1}$ |
| Cell concentration [1] ($K^+$) | $c_0$ | $mol.m^{-2}$ | 100 | $mol.m^{-3}$ |
| Hydrostatic pressure [2] | P | $N.m^{-1}$ | $10 - 100$ | $N.m^{-2}$ |
| Osmotic pressure | $\pi_0$ | $N.m^{-1}$ | $10^5$ | $N.m^{-2}$ |
| Tension [3] | $\gamma$ | $N$ | $5 \times 10^{-4}$ | $N.m^{-1}$ |
| Contact angle | $\theta$ | - | $\frac{\pi}{3}$ | - |
| Contact tension | $\gamma_c$ | $N$ | $\sim \gamma$ | $N.m^{-1}$ |
| Intercellular space width [4] | $e_0$ | $nm$ | 50 | $nm$ |
| Bridge length | $\ell$ | $\mu m$ | $0.1 - 10$ | $\mu m$ |
| Typical lumen length | $L_0$ | $\mu m$ | $0.1 - 10$ | $\mu m$ |
| Total length | $\mathcal{L}_0$ | $\mu m$ | $10 - 100$ | $\mu m$ |
| Water viscosity | $\eta$ | $N.s.m^{-1}$ | $10^{-3}$ | $N.s.m^{-2}$ |
| Water permeability [5] | $\lambda_v$ | $m^2.s^{-1}.N^{-1}$ | $7.32 \times 10^{-13}$ | $m^3.s^{-1}.N^{-1}$ |
| Solute permeability coefficient [6] | $\lambda_s$ | $mol^2.N^{-1}.s^{-1}.m^{-2}$ | $10^{-8}$ | $mol^2.N^{-1}.s^{-1}.m^{-3}$ |
| Solute diffusion constant (KCl) [7] | $D$ | $m^2.s^{-1}$ | $2.10^{-9}$ | $m^2.s^{-1}$ |
| Laplace/Osmotic pressures ratio | $\epsilon = \frac{\gamma \sin\theta}{L_0 \Pi_0}$ | - | $10^{-2} - 10^{-3}$ | – |
| Effective hydrodynamic friction | $\kappa_v = \frac{e_0^3}{12\eta}$ | $m^4.N^{-1}.s^{-1}$ | $1.04 \times 10^{-20}$ | $m^5.N^{-1}.s^{-1}$ |
| Pressure screening length | $\xi_v$ | $\mu m$ | 84 | $\mu m$ |
| Concentration screening length | $\xi_s$ | $\mu m$ | 14 | $\mu m$ |
| Active pumping flux | $j^a$ | $mol.s^{-1}.m^{-1}$ | $2.47 \times 10^{-10}$ | $mol.s^{-1}.m^{-2}$ |
| Solute time | $\tau_s$ | s | 2 | s |
| Water time | $\tau_v$ | s | 7 | s |
| Diffusion time | $\tau_D = \frac{\ell_0^2}{D}$ | s | $10^{-2}$ | s |

Table 1: Symbols, values and units.
